# Mouse fertilization triggers a conserved transcription program in one-cell embryos

**DOI:** 10.1101/2020.09.15.298018

**Authors:** Maki Asami, Brian Y. H. Lam, Martin Hoffmann, Toru Suzuki, Xin Lu, Matthew D. VerMilyea, Naoko Yoshida, Marcella K. Ma, Kara Rainbow, Stefanie Braun, Nina Patwary, Giles S. H. Yeo, Christoph A. Klein, Anthony C. F. Perry

## Abstract

Following fertilization, the new embryo reprograms parental genomes to begin transcription (embryonic genome activation, EGA). EGA is indispensable for development, but its dynamics, profile or when it initiates in vertebrates are unknown. We here characterize the onset of transcription in mouse one-cell embryos. Precise embryo staging eliminated noise to reveal a cascading program of *de novo* transcription initiating within six hours of fertilization. This immediate EGA (iEGA) utilized canonical promoters, produced spliced transcripts, was distinctive and predominantly driven by the maternal genome. Expression represented pathways not only associated with embryo development but with cancer. In human one-cell embryos, hundreds of genes were up-regulated days earlier than thought, with conservation to mouse iEGA. These findings provide a functional basis for epigenetic analysis in early-stage embryos and illuminate networks governing totipotency and other cell-fate transitions.

**One Sentence Summary:** Fertilization instates transcription in mouse and human one-cell embryos far sooner than thought and is programmed.

---

It is not known precisely when mammalian EGA initiates after fertilization (*1*). To address this, we generated high-resolution transcriptome profiles by single-cell microarray and RNA-seq analyses of tightly-synchronous one-cell mouse embryos collected at 2 h intervals after fertilization. Embryo synchronization in this way minimized noise to reveal hitherto inaccessible information about gene expression at the onset of development. Embryos of different crosses (designated F2 and B6*cast*) developed efficiently *in vitro* (fig. S1A) and their transcriptomes exhibited time-point-dependent clustering (Fig. 1, A and B). Merged F2 and B6*cast* (F2-B6*cast*) RNA-seq series detected 3,989 differentially-expressed genes (DEGs) relative to mature metaphase II (mII) oocytes (FDR<0.05; Fig. 1C and fig. S1, B to D), including most (64.8%; *n*=667) DEGs and pathway trends shared by microarrays (fig. S1E). We subsequently focussed on F2-B6*cast* RNA-seq series.

**Fig. 1.**
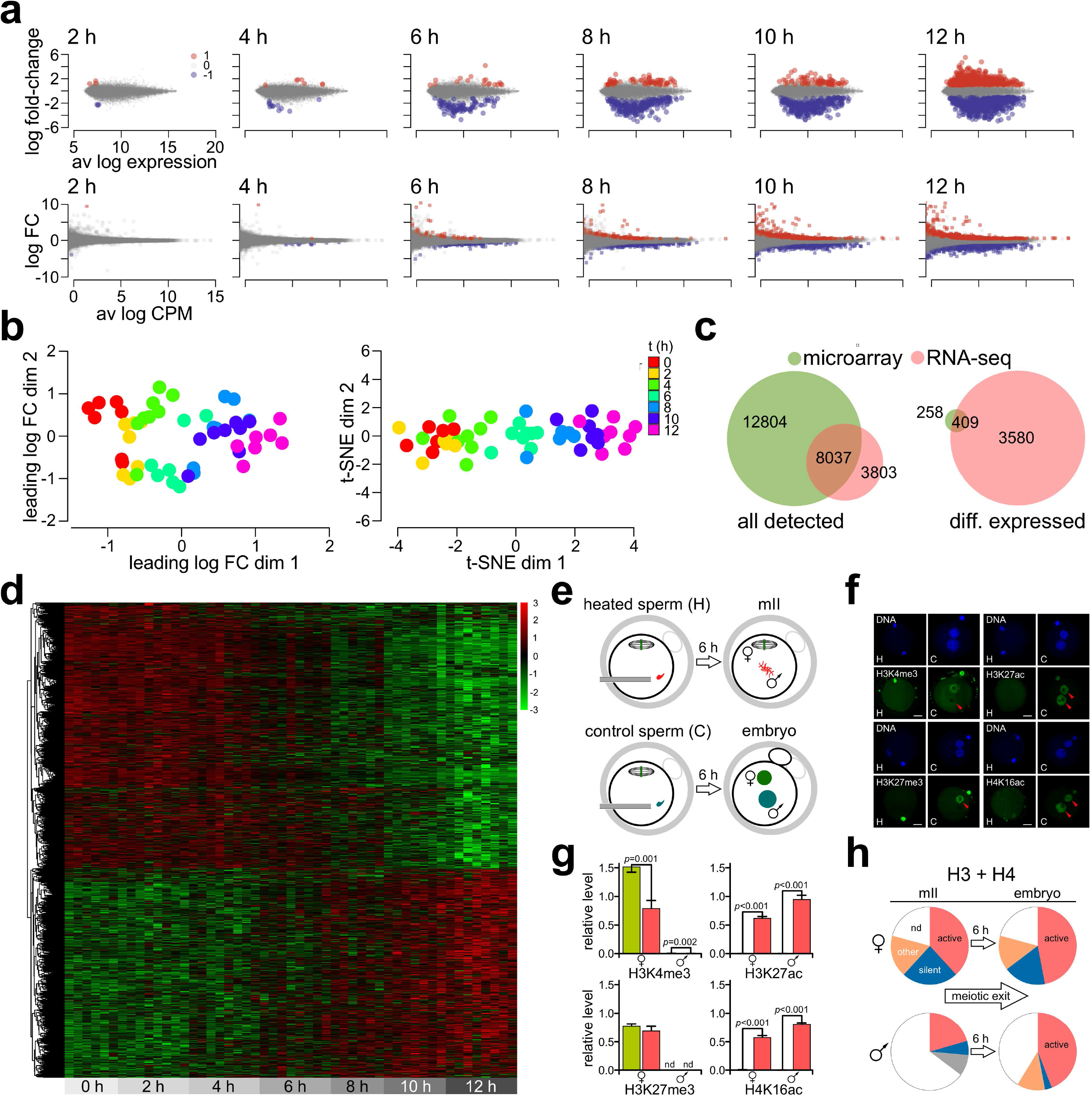
The dynamic transcriptome program of mouse one-cell embryos, 0-12 h. (**A**) Map-plots showing differentially-expressed genes relative to metaphase II (mII) oocytes from microarray (upper) and RNA-seq. (**B**) MDS analysis of microarray (left) and t-SNE analysis of RNA-seq data. (**C**) Relationships of microarray and RNA-seq data across the 12 h time-course, comparing all detected transcripts (left) and differentially-expressed transcripts (diff.). (**D**) Heatmap showing one-cell embryo gene expression changes (FDR<0.05). (**E**) Schematic of microinjection with control or heat-inactivated (heated) sperm. (**F**) Vertically-paired fluorescence images showing DNA (blue, Hoechst 33258) and histone modifications in biparental oocytes (mII) or embryos of (**E**). Arrowheads indicate paternal chromatin. Scale bar, 20 μm. (**G**) Quantification of images of (**F**) (5<*n*<9 oocytes or embryos each) as indicated at mII (green) or in embryos (red) after 6 h. Values show s.e.m. and isoparental chromatin labelling differences (*t*-test, *p*<0.05). (**H**) Pie charts summarizing histone modifications after 6 h with (right) or without meiotic exit.

Unsupervised cluster analysis of top F2-B6*cast* RNA-seq DEGs assigned each embryo to its correct relative position on the time-course (Fig. 1D). Of the 4,067 DEGs (FDR<0.05), 2,290 (56.3%) were down-regulated (fig. S2A), including maternal transcripts (*eg Mos, Plat* and *Gdf9*) (*2*-*4*). However, DEGs also reflected up-regulation that had clearly initiated by 6 h (Fig. 1D and fig. S1D). Genome activation within 12 h of fertilization (immediate EGA, iEGA) overlapped with EGA (*3, 5, 6*) but was distinctive (fig. S2), and genes responsive to the EGA regulator, Dux, were not up-regulated, further suggesting that EGA and iEGA are distinguishable (fig. S1F). Thus, mouse fertilization immediately triggers a signature program of gene expression, iEGA, in one-cell embryos.

EGA in non-mammalian vertebrates involves chromatin remodelling (*1, 7, 8*) leading us to monitor histone modifications (*n*=34) in mouse embryos and biparental mII oocytes containing both oocyte-and sperm-derived genomes (*9, 10*). Embryonic histone marks associated with gene activity increased in number by up to 2.5-fold compared to those in biparental oocytes (*p*=0.006; Fig. 1, E to H and fig. S3, A and B), suggesting that meiotic exit promotes active chromatin to enable iEGA.

Genes encoding detected iEGA transcripts were distributed uniformly throughout the genome (fig. S3C). Most were protein-coding, with annotated 5’-ends, implying that they mapped to canonical RNA polymerase II (Pol II) transcription start sites (fig. S3D), and some were refractory to the Pol II inhibitor, *α*-amanitin (fig. S4, A and B) as previously noted (*3*). Upstream regulator analysis (INGENUITY, FDR<0.05) of iEGA genes identified transcription factors (TFs) that included c-Myc and additional (*11*) cancer-associated TFs (Fig. 2A and fig. S5A). This lead us to investigate whether c-Myc might play a role in iEGA. c-Myc protein was present in mII oocytes, where it localized to spindles (*12*), and remained detectable in one-cell embryos (fig. S5, B to D). Genes encoding c-Myc-regulatory SUMO pathway components (*13*) were activated following fertilization (fig. S5E), consistent with an early embryonic role for SUMO. Moreover, inhibiting c-Myc from the onset of iEGA after fertilization lead to marked one-cell developmental arrest and blocked activation of candidate c-Myc-regulated iEGA genes (Fig. 2, B and C and fig. S5F). These findings suggest that iEGA gene expression is able to predict TFs that initiate embryonic transcription, that c-Myc is involved in iEGA and that iEGA shares regulatory features with cancer.

**Fig. 2.**
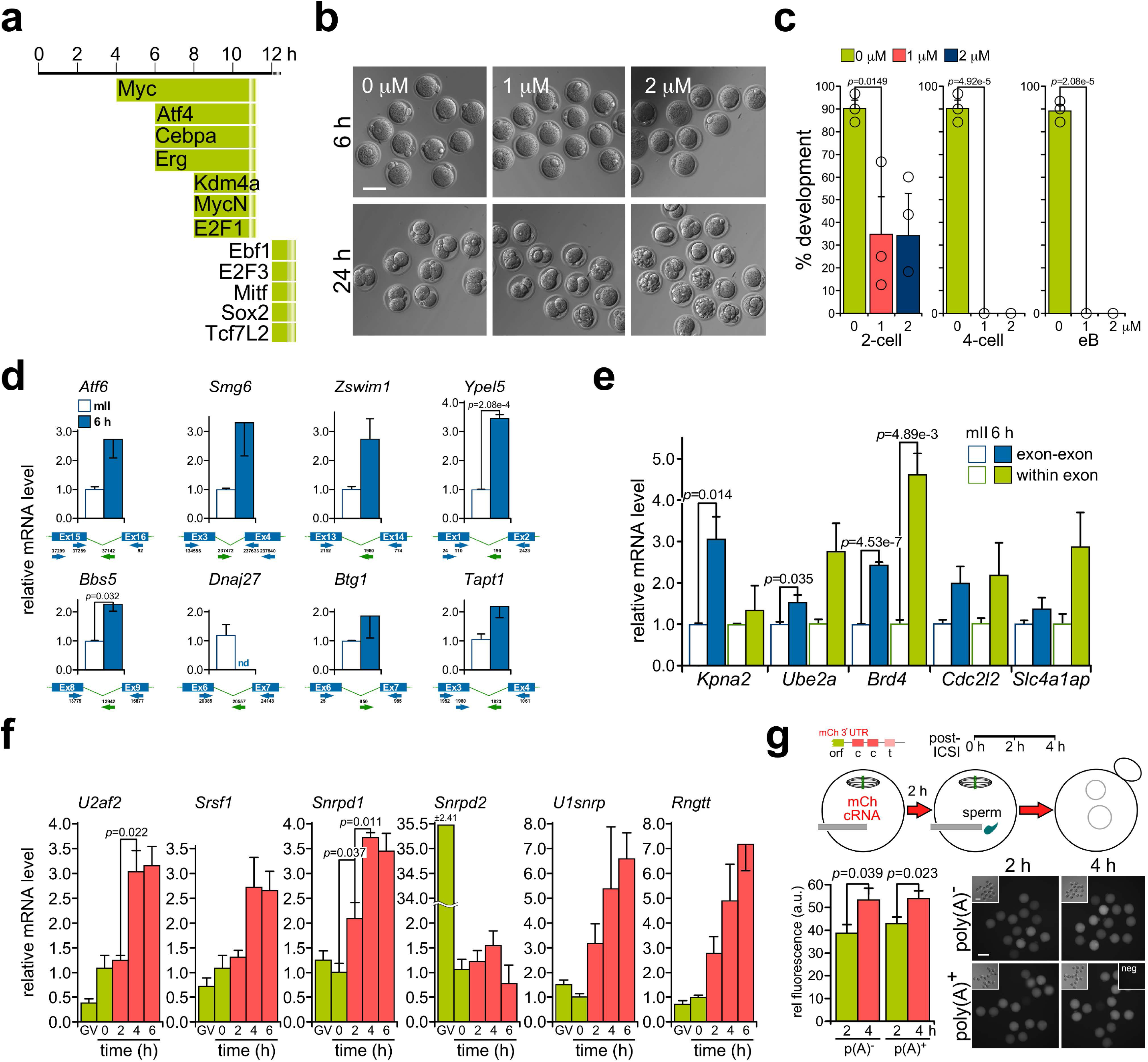
Transcript processing and translation immediately after fertilization. (**A**) Upstream transcription regulators inferred for each iEGA time-point (INGENUITY). (**B**) Hoffman micrographs of embryos produced by *in vitro* fertilization (IVF) and incubated in c-Myc inhibitor, 10058-F4 (0, 1 or 2 μM) after six (upper row) or 24 h. Scale bar, 100 μm. (**C**) Quantification of developmental rates (percentage of embryos surviving IVF) of embryos of (**B**) for *n*=3 independent biological replicates. 2-cell, two-cell embryo (24 h after IVF); 4-cell, four-cell embryo (∼48 h); eB, expanded blastocyst (∼96 h). (**D**) Histogram plots of transcript levels determined by qPCR using intron-flanking primers for random-primed cDNA derived from metaphase II (mII) oocytes (open bars) and embryos 6 h after *in vitro* fertilization. Intron-exon primer pairs gave products for genomic DNA but not cDNA. (**E**) qPCR analysis with intra-exonic primers and primers flanking exon-exon junctions. (**F**) Spliceosome component and guanylyltransferase (*Rngtt*) transcript levels determined by qPCR in germinal vesicle (GV) oocytes, mII oocytes (0) and one-cell embryos at the times shown (h) after sperm injection. (**G**) Injection of *mCherry* cRNA (mCh, top left: orf, mCherry open reading frame; c, cytoplasmic polyadenylation element; t, mRNA cleavage/polyadenylation signal). Fluorescence intensity quantification (lower left) at the times shown after injection of *mCherry* cRNA (0.6 ng/μl) polyadenylated *in vitro* (pA^+^) or not (pA^-^). Fluorescence micrographs show representative oocytes with corresponding bright field images (insets, upper left) and a non-injected control (neg, inset upper right). Bars, 100 μm. Values in (**C**-**G**) are ± s.e.m. Unpaired *t*-tests show *p*-values <0.05.

Most (2759/2811, 98.2%) iEGA transcripts exhibited canonical splicing (Fig. 2, D and E, and fig. S1, C and G) and genes for core spliceosomal components and the mRNA capping enzyme, guanylyltransferase (*Rngtt*) were up-regulated by 6 h (Fig. 2F). iEGA transcripts were processed independently of polyadenylation (fig. S6, A and B) (*4*) and some corresponded to protein level increases (fig. S6C); transcript interference *via* a splice-blocking morpholino could reduce corresponding protein levels within 6 h, supporting a nascent transcript contribution (fig. S6D). Oocytes coinjected with sperm plus *mCherry* cRNA expressed mCherry protein within 4 h whether or not the injected cRNA had been polyadenylated (Fig. 2G). Thus, iEGA is accompanied by translational competence.

The F2-B6*cast* time-course revealed a pathway succession fitting developmental processes (Fig. 3A and fig. S6E) (*14, 15*) and endogenous retrovirus (MuERV) *LTR, Pol* and *Gag* transcripts (*16*) were up-regulated at hundreds of loci from ∼8 h (Fig. 3B), showing that *MuERV* gene activation is part of iEGA. To discriminate between parental genomes in iEGA, we analysed unmerged B6*cast* (*Mus musculus domesticus* x *M. m. castaneus*) series transcriptomes. Clustering analysis separated B6*cast* embryos along the time-course (fig. S7A) and permitted parent-specific expression assignment. Informative transcript changes predominantly reflected maternal mRNA degradation (in 2226/4413 cases; logFC<0; Fig. 3, C and D) followed by maternal allelic gene expression (logFC>0; Fig. 3C). We investigated whether sperm-borne mRNA was introduced during removal of the major sperm nucleoprotein, protamine following fertilization. In agreement with a previous approximation (*9*), 50% of transgene encoded fluorescent protamine was removed by ∼30 min (fig. S7, B to F). However, starting the B6*cast* time-course at 20 min did not detect significant (FDR<0.1) levels of sperm-borne mRNA (*17*). These findings argue against the delivery of a major cargo of sperm-borne mRNA into oocytes and that iEGA predominantly involves maternal gene expression.

**Fig. 3.**
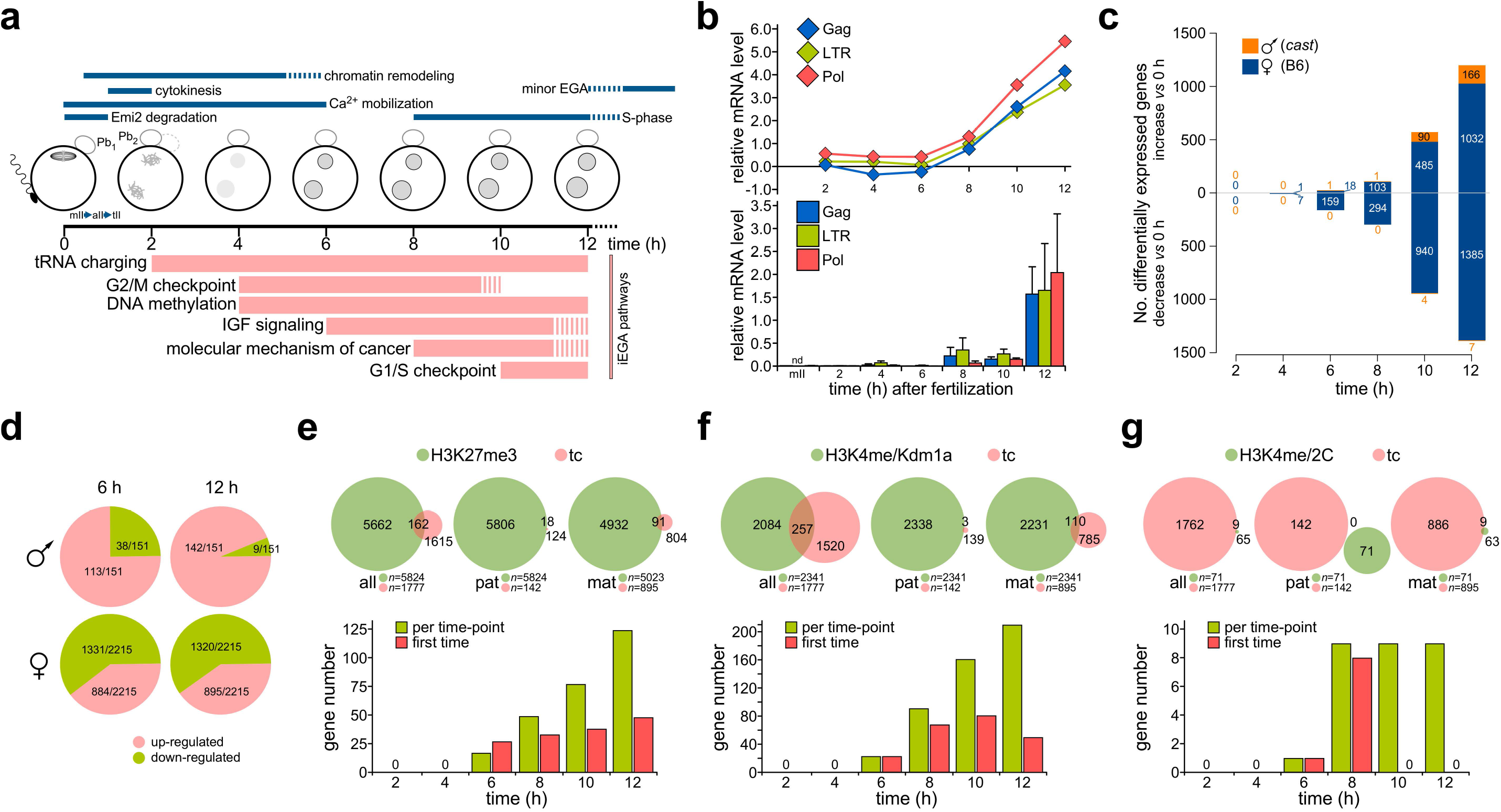
Mouse one-cell embryo transcriptome signature. (**A**) iEGA pathway succession and embryonic events in the first 12 h. Pb_1_, first polar body; Pb_2_, second polar body; mII, aII and tII, second meiotic prophase, metaphase, anaphase and telophase respectively. (**B**) Relative levels (±s.e.m.) of *MuERV* transcripts *LTR, Pol* and *Gag* (RNA-seq; FDR *p*-value ≤0.05) at the times shown post-fertilization, from RNA-seq (upper) and qPCR of pools of embryos generated by IVF (*n*=6 pools/time-point). (**C**) Histograms showing transcript level changes compared to mII oocytes (0 h) for maternal (B6) and paternal (*cast*) alleles in the B6*cast* series (FDR *p*-value <0.05). (**D**) Percentages of genes (RNA-seq; FDR *p*-value ≤0.05) from each parental genome at times indicated post-fertilization. (**E**) Venn diagrams (upper) of up-regulated iEGA genes in F2-B6*cast* (all) and B6*cast* paternal (pat) and maternal (mat) RNA-seq datasets, as they map to promoters marked with H3K27me3 in mII oocytes (*18*). Histograms for F2-B6*cast* data show cumulative up-regulated gene numbers and numbers up-regulated for the first time. (**F**) As per (**E**), showing overlaps with H3K4me2 promoter occupancy in sperm altered by exposure to transgene expression of the H3K4me demethylase, Kdm1a (*20*). (**G**) As per (**E**), showing overlaps with genes differentially expressed in two-cell embryos (2C) following fertilization by transgenic *Kdm1a* sperm (*20*).

We next asked whether regulatory epigenetic marks were present at iEGA genes. Only one iEGA gene contained a DNA methylation imprint (*Flt1*; *p*<0.05), but some iEGA promoters contained the repressive mark, H3K27me3, present at the same promoter in sperm (*18*) or mII oocytes (*19*), or the active one, H3K4me2 in sperm (*20*) (*p*<0.05; Fig. 3, E and F, and Table S1). Further data mining revealed 17,168 promoter regions associated with the active mark, H3K4me3 in the sperm genome (FDR<0.05) (*21*), of which 1,116 corresponded to iEGA genes (62.8% of the total; *n*=1,777; FDR<0.05). Only 95 (5.3%) iEGA promoter regions were present in the set of promoters bivalently marked in mouse primordial germ cells (*n*=3,498; FDR<0.05) (*22*). Assuming that the marks are preserved in the germline (*22*), this indicates that iEGA occurs independently of bivalently-marked gamete promoters.

Human EGA is thought to initiate between four-and eight-cell stages (*23*-*25*), leading us to perform high-resolution single-cell RNA-seq of human mII oocytes (*n*=12) from six donors aged 23-31, and one-cell embryos (*n*=12) from four unrelated couples. Unsupervised clustering analysis respectively placed oocyte and embryo transcriptomes into discrete groups (Fig. 4, A to C, and fig. S7G). In agreement with mouse data, most DEGs in human one-cell embryos (1,121 out of 1,995) were down-regulated, with 874 up-regulated (FDR<0.05; logFC>0) including orthologs of 132 (15%) mouse iEGA genes (Fig. 4, D and E). Human DEGs included 16 down-and 29 up-regulated *hERV* loci (Table S2). Predicted pathway terms (fig. S8) were shared with mouse (Fig. 4F) and included transcriptional networks regulated by MYC and MYCN (Fig. 4G and fig. S9). Transcripts for putative TF drivers of human cleavage-stage EGA (*eg DUX4, OCT4* and *LEUTX*) (*24*-*27*) were not up-regulated. Thus, mouse and human iEGA overlap, suggesting that iEGA is broadly conserved in mammals.

**Fig. 4.**
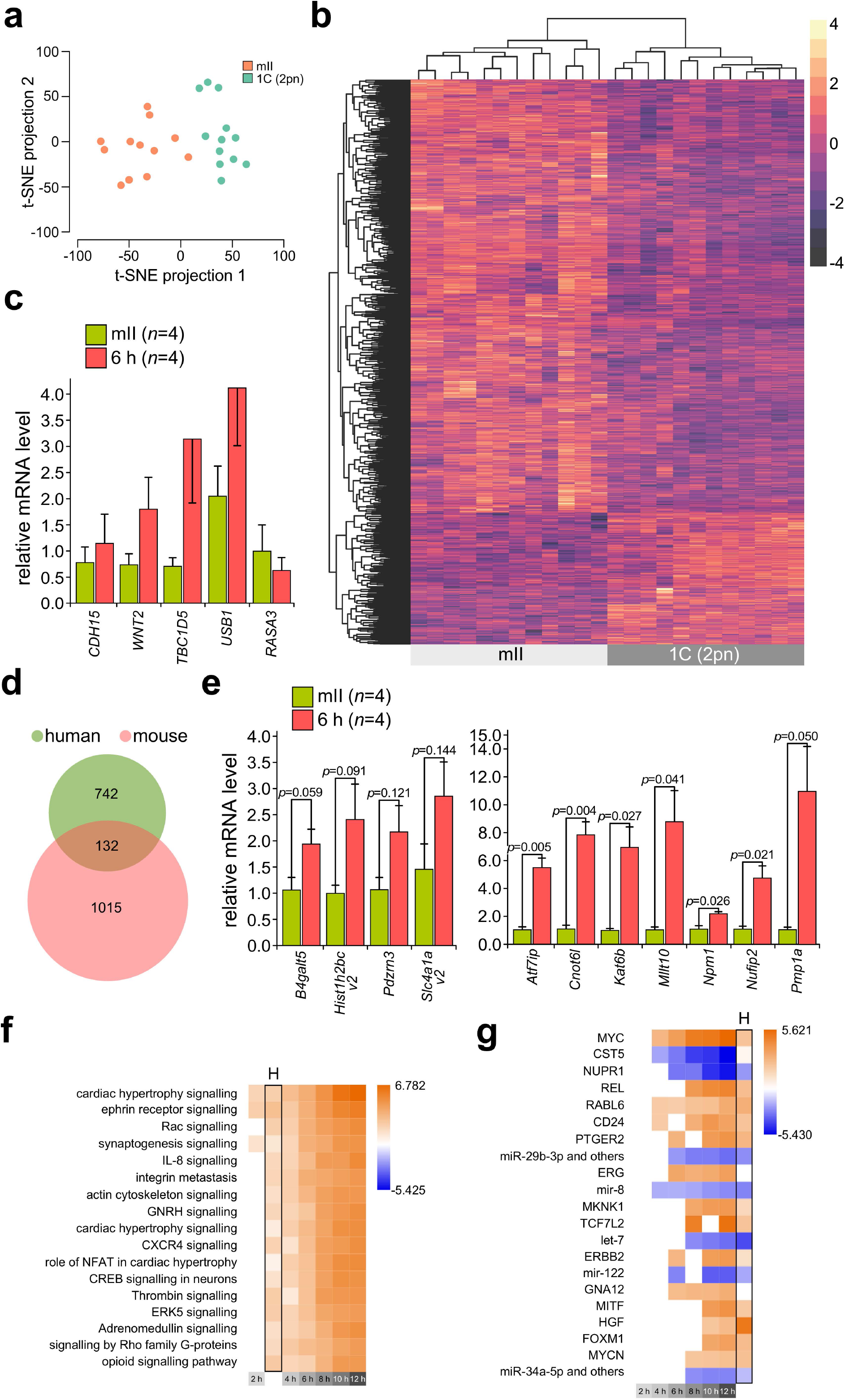
Human embryonic transcription initiates at the one-cell stage. (**A**) t-SNE analysis of RNA-seq data for human metaphase II (mII) oocytes (*n*=12) and one-cell embryos (*n*=12) from different donors. (**B**) Heatmap showing changes in transcript levels (FDR *p*-value <0.10) in human bipronuclear one-cell (1C, 2pn) embryos (*n*=12) and mII oocytes (mII) (*n*=12) of (**A**). (**C**) Histograms of qPCR for single human one-cell embryo (*n*=4 separate embryos each) transcripts up-regulated from RNA-seq. Different embryos were used than for (**A**). Values are ± s.e.m. and normalized against mII oocytes (∼1.0). (**D**) Venn Diagram showing up-regulated gene overlap between human (FDR<5%, logFC>0) and mouse (full time-course FDR<5%, slope >0) one-cell embryos. (**E**) Histograms for transcript levels (qPCR) of mouse orthologs of transcripts up-regulated in human one-cell embryos. Values are ± s.e.m., normalized against mII oocytes (∼1.0). *p*-values (unpaired *t*-test) are indicated for pair-wise comparisons. (**F**) Pathway analysis of up-regulated genes in human (H) and mouse one cell embryos. (**G**) Transcription regulators suggested for human (H) and mouse one-cell embryos indicated by INGENUITY analysis.

This work reports an iEGA program that anticipates key events during the establishment of embryonic totipotency and is predominantly maternally-driven (Fig. 3A) (*14, 15, 28*). Maternal genome predominance is possibly an evolutionary corollary of parthenogenetic developmental potential in some non-mammalian vertebrates (*29*), prevented in mammals by imprinting (*30*). The distinctive stereotypical nature of iEGA and its association with the extreme cell fate change, from gamete death to the establishment of totipotency (*28*), hint that it is archetypal: many potency changes may ‘borrow’ from it. It is therefore possible that disease states (*eg* some cancers) involving cell fate changes arise due to ectopic expression of the iEGA program, and that iEGA is predictive of disease mechanisms. Unlocking iEGA in its natural embryonic context thus promises to open new doors to regulatory (*eg* epigenetic) processes that are required for embryogenesis but cause disease if they become dysregulated in adulthood.

## Acknowledgements

We thank Animal Facility support staff, J. Whitney for human embryo curation and C. Tickle for comments during manuscript preparation.

## Author contributions

A.C.F.P. conceived the core idea, and (with M.A.) molecular and imaging experiments, performed by M.A., M.D.V. and N.Y. Mouse embryology was by M.A, T.S. and A.C.F.P. Human embryo preparation was by M.D.V. and S.B. Single-cell whole transcriptome amplification was performed by N.P. (microarray) and B.L., K.R. and M.K.M. (RNA-seq). C.A.K. designed microarray analyses, performed and evaluated by M.H., X.L. and C.A.K. B.L. designed RNA-seq analyses, performed and evaluated by B.L., M.M. and G.S.H.Y. Data analysis was by M.A., B.L., G.S.H.Y. and A.C.F.P. A.C.F.P. wrote the manuscript.

## Data and materials availability

All data are available in the main text or supplementary materials. Gene Expression Omnibus accession numbers for expression data are GSE64650 (microarray) and GSE157834 (RNA-seq).

## Supplementary Information

### Materials and Methods

#### Animal care

Experiments involving animals were performed in accordance with local and national statutes including the University of Bath Animal Welfare Ethical Review Body and complied with the UK Animals (Scientific Procedures) Act, 1986 and its embodiments. The study did not involve wild animals.

#### Collection and culture of mouse oocytes

Oocytes were from a *Mus musculus domesticus* C57BL/6 (B6) x DBA/2 F1 hybrid and *M. m. castaneus* (*cast*), and embryos from B6 x DBA/2 F2 hybrids (F2), or B6 x *cast* (B6*cast*) crosses. Oviductal metaphase II (mII) oocyte complexes were typically collected in M2 medium (EMD Millipore, UK)^31^ from 8-12-week-old C57BL/6 or B6D2F_1_ females (produced by crossing C57BL/6 females with DBA/2 males in-house or otherwise supplied by Charles River; L’Arbresle, France) 12 to 15 h after standard superovulation by serial injection of equine and human chorionic gonadotropin (PMSG and hCG). Complexes were then either used in IVF (below), or cumulus cells were removed by hyaluronidase treatment and after multiple washing in M2 medium, denuded oocytes incubated in kalium simplex optimized medium^32^ (KSOM; Millipore) under mineral oil in humidified 5% CO_2_ (v/v air) at 37°C, until required^33^.

#### Sperm preparation and microinjection (ICSI)

Preparation of cauda epididymidal sperm from 8-to 12-week-old *Mus musculus musculus* B6D2F_1_ or *Mus musculus castaneus* (*cast*) males for ICSI was essentially as previously described^33,34^. Injection was completed in ± ≤2.5 min for each timepoint. The B6*cast* series was generated by injecting *cast* sperm into C57BL/6 oocytes. To prepare sperm, they were triturated for 45 sec in nuclear isolation medium (NIM; 125 mM KCl, 2.6 mM NaCl, 7.8 mM Na_2_HPO_4_, 1.4 mM KH_2_PO_4_, 3.0 mM EDTA; pH 7.0) containing 1.0% (w/v) 3-[(3-cholamidopropyl)dimethylammonio]-1-propanesulfonate (CHAPS) at room temperature (25°C). Sperm were washed twice in NIM and pelleted at ambient temperature; head-tail detachment was enhanced by trituration during pellet resuspension. Finally, sperm were resuspended in ice-cold NIM (∼0.5 ml per epididymis equivalent) and stored at 4°C for up to 3 h until required, but typically injected immediately after preparation. This gentle protocol removes sperm membrane and other (*eg* acrosomal) components that do not enter the oocyte during fertilization; the sperm support normal, healthy full-term development^17,34^.

Immediately before microinjection, ∼50 μl of each sperm suspension was mixed with 20 μl of polyvinylpyrrolidone (PVP, average *M*_r_ ≈ 360,000; Sigma-Aldrich) solution (15% [w/v]) and sperm injected (ICSI) within ∼60 min of PVP mixing into oocytes in a droplet of M2 as described^33,34^. After a brief recovery (∼5 min), injected oocytes were transferred to KSOM under mineral oil equilibrated in humidified 5% CO_2_ (v/v air) at 37°C and cultured until required.

Previous studies on early embryonic transcription in the mouse^3,24,25,35-37^ have tended to generate embryos by natural mating following superovulation, with the timing of embryo collection relative to the time of hCG injection rather than fertilization, resulting in poor, or no developmental synchrony and the loss of critical information to variation between individual embryos. Natural mating further precludes synchronization because its precise timing is difficult to ascertain. Even were the time of coitus known, in the mouse, the duration of sperm passage through the uterotubal junction to the oviduct varies^38^, sperm fusion with all oocytes occurs over a period of ∼2.5 h^39^, and oocytes remain in a fertilizable state for at least 12 h post-ovulation^40,41^. This contrasts with the sperm injection (ICSI) protocol used here, which can generate synchronized cohorts (∼10 mII oocytes can readily be injected in 5 min) of developmentally competent embryos with precision^33^. There are no significant developmental differences between ICSI, *in vitro* fertilization (IVF) and natural mating^42^ and in some cases, ICSI may even be developmentally superior^43^. ICSI is currently the main method of assisted human reproduction worldwide, with 66.5% of total nondonor aspiration cycles worldwide^44^ and studies on mouse ICSI are highly relevant to clinical practice.

#### Mouse *in vitro* fertilization (IVF)

Where appropriate, embryos were generated by standard B6D2F_1_ × B6D2F_1_ IVF. Sperm were collected from mature males by epididymidal puncture followed by dispersal for 5 min in pre-warmed human tubal fluid (HTF; Millipore) in humidified CO_2_ (5% [v/v] in air) at 37°C. Of the 400 μl dispersal droplet, 10 μl was transferred to a fresh fertilization dish containing 200 μl HTF and incubation continued for 1 h before placing ∼30 cumulus oophorous complexes freshly-isolated from superovulated 8-week-old B6D2F_1_ females and incubating in the CO_2_ incubator at 37°C. The resultant embryos were washed in fresh HTF and dead and clearly unfertilized oocytes removed. Embryos were then washed 5x in KSOM and incubated until required in KSOM droplets equilibrated under mineral oil in humidified 5% CO_2_ (v/v air) at 37°C.

#### Mouse embryo culture

Embryos were typically cultured in KSOM droplets equilibrated under mineral oil in humidified 5% CO_2_ (v/v air) at 37°C as previously described^10,33,41,48-52,56^. To inhibit the transactivation of c-Myc target gene expression, embryos produced by IVF 2 h after sperm-oocyte mixing, were washed and transferred to KSOM supplemented with the inhibitor, 5-[(4-Ethylphenyl)methylene]-2-thioxo-4-thiazolidinone (10058-F4; Sigma-Aldrich) and incubation continued. Dimethylsulphoxide (DMSO) was used to solubilize 10058-F4 (with 10058-F4 at working concentrations of 1.0 to 3.0 μM) and was included in media without 10058-F4 for negative controls.

#### Human metaphase II oocytes and zygotes (one-cell embryos)

Patients underwent ovarian stimulation according to guidelines of each clinic, where protocols included agonist luteal phase and antagonist suppression. On the day of mII oocyte retrieval (day 0), oocytes were either cryopreserved by a slow freeze method using propanediol (PROH)^45^ or used to produce embryos by *in vitro* fertilization or ICSI. Immediately after fertilization assessment, morphologically normal bipronuclear (2pn) zygotes were cryopreserved using dimethylsulfoxide^46^ and stored under liquid nitrogen as appropriate. Some sibling embryos gave rise to children, further attesting to their healthy status. When required, cryopreserved oocytes and zygotes were thawed by rapid warming using a Vit-Warm Kit (FUJI Irvine Scientific, USA) according to the recommended protocol, and viability confirmed. All mII oocytes and zygotes were washed in protein-free multi-purpose handling medium (FUJI Irvine Scientific, USA) and each placed in a 0.8 ml PCR tube containing 1x single-cell lysis buffer supplemented with RNase inhibitor (Takara Clontech, USA). Oocyte and one-cell embryo donor groups did not overlap; mII oocytes and zygotes came from different individuals. There were six mII oocyte donors (aged 23, 24, 24, 25, 27 and 31 years); five were Caucasian and one African American/Hispanic. There were four zygote donor couples; for two the male and female ages were respectively 36 and 38 and 40 and 50 (data are unavailable for the other two couples) and three of the couples were Caucasian, with one Asian couple.

#### Immunocytochemistry

Oocytes, embryos and cultured cells were fixed in 4% (w/v) paraformaldehyde and either processed immediately or stored at 4°C for up to 2 weeks until required. Fixed cells were permeabilized by incubation in PBS supplemented with 0.5% (v/v) triton X-100 and 0.1% (w/v) BSA for 30 min at 37°C, followed by blocking in PBS supplemented with 3% (v/v) normal goat serum and 0.1% (w/v) BSA for 30 min at room temperature. Labeling was by incubating samples overnight at 4°C in primary antibody followed by a incubation for 1 h at 37°C with the appropriate secondary antibody (1:250 [v/v]; Life Technologies Ltd., UK) conjugated to Alexa 488, Alexa 594 and Alexa 647. DNA was stained by incubating samples at 37°C for 20 min in propidium iodide (1:200 [v/v]; Sigma, USA) or Hoechst 33342 (1:1000 [v/v]; Sigma). Chromatin epitopes in mII oocytes and embryos are accessible to their cognate antibodies^10^ and most (27/34) here resided on solvent-exposed N-terminal histone tails^47^, although in 28/34 cases (82.3%), samples contained in-built positive controls in which one or both parental chromatin sets stained within a single given cell. Only one situation corresponded to an epitope (the core modification, H4K91ac) that was unrecognised in both parental alleles at mII, and both in one-cell embryos. Antibodies with no reactivity in either mII oocytes or embryos were excluded from the analysis.

#### Direct fluorescence imaging and analysis

Differential interference contrast microscopy (DIC) and epifluorescence imaging have been described previously (*41, 48*-*50*). Images of live oocytes or embryos following cRNA injection were captured on an Olympus IX71 equipped with an Andro Zyla sCMOS camera and OptoLED illumination system (Cairn Research Ltd., UK) and processed using Metamorph software (Molecular Devices, LLC, USA). Excitation at 587 nm in combination with an ET-mCherry filter system was used for mCherry fluorescence detection and at 484 nm with an ET-EYFP filter system to detect Venus epifluorescence. Confocal images were obtained using LaserSharp 2000 6.0 Build 846 software on an Eclipse E600 (Nikon, Japan) microscope equipped with a Radiance 2100 laser scanning system (BioRad, USA; LSM Technical Service, UK). Image J (http://rsbweb.nih.gov/ij) was used in image data analysis. The parental provenance of pronuclei in mouse one-cell embryos was assigned according to size and position; the female pronucleus is consistently smaller and closer to the second polar body than the paternal pronucleus.

#### Preparation and injection of cRNA

5’-capped cRNA was synthesized *in vitro* from linearized plasmid template DNA in a T7 mScript(tm) Standard mRNA Production System (Cellscript, USA) according to the recommendations of the manufacturer, as previously described^48,51,52^. Where appropriate, the polyadenylation step following synthesis was omitted (Fig. 2h). cRNA was dissolved in nuclease-free water, quantified on a Nanophotometer and stored in aliquots at ^-^80°C until required. cRNA solutions were diluted as appropriate with sterile water and injected (typically at concentrations of 0.01 to 1 µg/µl) within 1 h of thawing *via* a piezo-actuated micropipette into mII oocytes or embryos in M2 medium.

#### Ratiometric PCR (qPCR)

For ratiometric transcript quantification by PCR (qPCR), embryos were produced either by IVF or ICSI and incubated in humidified 5% CO_2_ (v/v air) at 37°C until required. At the appropriate time post-injection, embryos were examined to confirm morphology (*eg* the presence of a second polar body and two pronuclei), transferred to 200 μl Isogen (Nippon Gene, Japan) containing 10 ng tRNA (Hoffmann-La Roche Ltd., Basel, Ch) and either used directly or flash-frozen in liquid nitrogen and stored at ^-^80°C until required. Synthesis of cDNA from total RNA was primed with random 8-mers plus oligo(dT)_20_ (each at 30 μM) in a 21 μl reaction volume containing 200 U SuperScript III reverse transcriptase (Invitrogen). For other qPCR, total RNA was extracted by transferring 5∼10 oocytes or embryos in a minimal volume (<0.5 μl) into 1 μl 0.1% (w/v) Sarkosyl (Teknova, Hollister, CA) containing 10 ng tRNA (Hoffmann-La Roche Ltd., Basel, Ch), heated at 65°C for 5 min and used to program cDNA synthesis primed with random 6-or 8-mers, with or without oligo(dT)_20_ (each at 30 μM) in a 21 μl reaction volume containing 200 U SuperScript IV reverse transcriptase (Thermo Fisher Scientific, UK). qPCR reactions were performed in a QuantStudio 7 (Thermo Fisher Scientific, UK) or ABI 7500 Real Time PCR System (Applied Biosystems, CA) in reactions (20 μl) containing 1-2 μl template cDNA, forward and reverse primers (100 nM each) and 12.5 μl of Power SYBR (ABI), using the parameters: 10 min at 95°C, followed by up to 45 cycles of (15 sec at 95°C, 1 min at 58°C and 35 sec at 72°C). Each experiment was performed with biological triplicates collected on at least two days and included technical duplicates of each sample. Primer sets (Hokkaido System Science, Japan or Eurofins MWG Operon, Germany) were non-dimerizing under the conditions employed. Reactions lacking input cDNA were used to verify absence of contamination in cocktail components. Steady state transcript levels were normalized with respect to the internal reference, *H3f3a* based on work by ourselves and others showing that *H3f3a* is robustly expressed in mouse oocytes and preimplantation embryo^41,48,53-56^. The CT value for *H3f3a* corrected for cell number (CT+log2 embryo cell number) is constant during preimplantation stages, giving a mean value of 29.56±0.23 (for 25≤*n*≤29 independent replicates). Statistical differences between pairs of data sets were analyzed by a two-tailed unpaired *t*-test and *p*-values ≤0.05 considered statistically significant unless stated otherwise.

#### Microarray analysis

For time-course transcriptomic microarray analysis, oocyte and embryo transcriptomes were prepared as for RNA-seq and preparation and whole transcriptome amplification fidelity assessed as previously described^48,56-58^. Gene expression data were quality-assessed by inspection of chip raw images and gene expression frequency distributions. Only high quality data were approved for further bioinformatic analysis producing 51 samples. Raw gene expression data were background-corrected (limma R-package, normexp method)^59^ normalized by quantile normalization. Technically replicated probes (identical Agilent IDs) were replaced by their median per sample. The original standard deviation was 2.07. Technically replicated probes (identical Agilent IDs) were replaced by their median per sample. For clustering and functional annotation, probes targeting the same gene were disambiguated by retaining only the probe with the lowest *p*-value. This reduced the 41,000 non-control probes to 29,078, of which 21,391 were annotated by proper gene symbols. The number of exons was retrieved from UCSC known genes (mm9) by matching Agilent probe sequence locations to UCSC transcript coordinates. If different transcripts for the same probe sequence were available the number of exons was averaged across these transcripts.

#### Mouse single-cell RNA sequencing

Single-cell oocytes and embryos 2, 4, 6, 8, 10 and 12 h post-ICSI were lysed in 1x single-cell lysis buffer containing RNase inhibitor (Takara Clontech, USA). RNA from lysates was subjected to direct reverse transcription with template-switching oligo and random hexamers, followed by PCR amplification, Illumina sequencing library generation and ribosomal RNA removal (all using Clontech SMARTer Total RNA-Seq Kit Pico Input V1). The libraries were then combined at equal molar concentrations and sequenced both ends for 100 base pairs (PE100) on an Illumina HiSeq 4000 instrument.

#### Human single-cell RNA sequencing

The protocol we used was similar to the one for mouse single-cell RNA sequencing. Human mII oocytes and embryos were lysed using 1x single-cell lysis buffer containing RNase inhibitor. Lysates were then used to generate RNA sequencing libraries using Clontech SMARTer Total RNA-Seq Kit Pico Input (V2), which were sequenced from both ends for 100bp (PE100) using an Illumina HiSeq 4000 instrument.

#### Sequencing bioinformatics

Raw sequence reads (fastq) from mouse oocytes and embryos were trimmed using the fastx_trimmer command from fastx-Toolkit (version 0.0.13; respectively 4 nt from 5’ and 15 nt from 3’ ends), and sequencing adapter sequences were removed by Cutadapt (1.7.1). Mapping of the trimmed reads was by Tophat (2.0.11) using the GRCm38 genome and Ensembl 54 reference transcriptome. After sequence alignment, the transcriptome was remodelled *via de novo* re-assembly of transcripts based on empirical data from mapped reads and incorporation into the original reference (Cufflinks 2.2.1). Gene-level counts based on the updated transcriptome were then performed using ht-seq-count (0.6.1p1) and transcript level counts *via* cuffnorm command from Cufflinks (2.2.1).

For the parent-of-origin study, gffread utility from Tophat (2.0.11) was used to generate the reference fasta for the transcriptomes of C57/BL6 and Cast/EiJ based on their corresponding reference genomes (GRCm38, Ensembl 70) and remodelled transcriptome data as described above. The transcriptomes were merged to form a single ‘F1’ reference transcriptome for mapping of trimmed sequence reads using Bowtie 1.1.0. Mapped reads were subsequently analysed using MMSEQ 1.0.9 to estimate the transcript level and aggregated gene level abundance originating from the genome of each strain. Abundance tables were generated using mmseq.R R script accompanied by the MMSEQ package for downstream analyses.

Merged F2 and B6*cast* RNA-seq series (*n*=52 samples, 6≤*n*≤9 per time-point; median 29 million reads per sample) detected 11,840 unique genes with >1 count per million reads. For human sequencing data, reads were mapped onto the Human GRCh38 genome and Ensembl 92 transcriptome using STAR (2.5.0a). Post-alignment reads were then fed into Stringtie (1.3.6) to remodel the transcriptome to assemble and incorporate novel transcripts, analogous to mouse RNA-seq analysis. The gene-level count was performed using htseq-count 0.6.1p1 and the transcriptome remodelled using Stringtie for downstream differential expression analysis. This analysis yielded a median of 67 million reads per sample.

#### Differential expression and pathway analysis

Raw gene-level counts were analysed using edgeR. A generalised linear model (GLM) was applied to determine dispersion, and likelihood-ratio test was used to detect differential gene expression over the time-course. For estimated counts downstream of Cufflinks and MMSEQ, data was normalised and mean variance relationships determined using limma-voom, followed by empirical Bayesian moderated F-statistic or *t*-test (eBayes) as appropriate. For the microarray dataset, raw gene expression values were background-corrected and normalized by quantile normalization using the Limma R package. As with RNA sequencing analysis, a GLM was fitted and eBayes used to detect differential gene expression over the time series. Pathway and upstream regulator analysis were performed using Qiagen Ingenuity Pathway Analysis software.

#### Sperm H3K4me3 ChIP-seq analysis

A sperm H3K4me3 ChIP-seq dataset was downloaded from Gene Expression Omnibus (Accession GSE42629; [Ref. 21]). Raw fastq reads for H3K4me3-bound DNA (GSM1046833, ChIP) and sonicated genomic DNA from sperm (GSM1046836, input control) were aligned to the mouse GRCm38 genome using BWA MEM (v0.7.12). Mapped reads were sorted by their genomic coordinates and used as input for peak detection using MACS2 (Ref. 60), which performed PCR de-duplication, library normalization, ChIP peak modelling and calling by comparing reads from the H3K4me3 bound library to background controls (sperm genomic DNA). FDR<0.05 and peak fold-enrichment >5 were used to filter peak calls (total 44,209 peaks spanning a total of 62 Mb, or ∼1.78% of the genome). Filtered peaks were annotated using ChIPSeeker R package with Ensembl mouse reference gene model GTF version 100. Of the peaks, 28,190 (63.8%) were labelled within the promotor (±3 kb from the transcriptional start site [TSS]) and 5’ untranslated regions (UTRs). Only peaks within the proximal promotor region (*ie* 1.5 kb upstream or downstream of the TSS) were used for downstream comparisons.

#### Data availability

Source Data are provided for figures. Microarray and RNA-seq data have been deposited into Gene Expression Omnibus (GEO).

#### Statistics and reproducibility

Statistical differences between pairs of data sets were analysed by two-tailed unpaired *t*-tests. Values of *p*<0.05 were considered statistically significant unless stated otherwise.

### Supplementary figure legend

**Figure S1.**
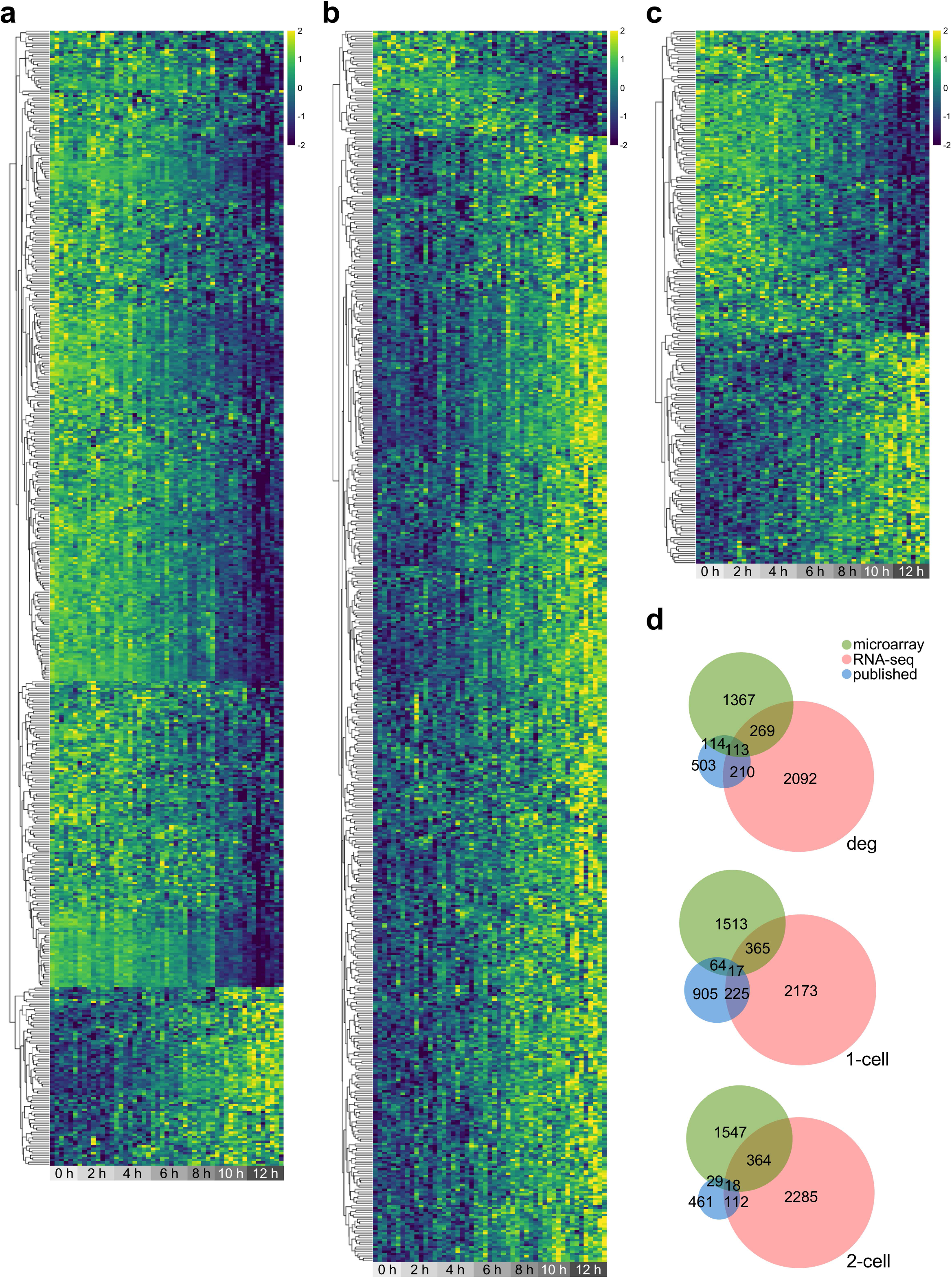
Transcript level overlaps with published data. **a**, Heatmap showing the overlap between transcripts exhibiting expression changes across the F2-B6*cast* 12 h time-course (FDR *p*-value <0.05) and previously-reported maternal genes^61^. Most genes map to early time-points, consistent with rapid transcript degradation. **b**, Heatmap as per (**a**), but showing the overlap with minor EGA genes as previously defined^61^. **c**, Heatmap as per (**b**), but showing the overlap with major EGA genes as previously defined^61^. **d**, Venn diagram showing numbers of overlapping genes whose transcript levels change throughout the time-course (deg) or were previously reported in one-cell (1C) or two-cell (2C) embryos (published)^61^.

